# Auto-downregulation of the florigen FT production prevents precocious flowering in plants

**DOI:** 10.1101/2025.04.24.650415

**Authors:** Shu Tian, Xiao Luo, Bowen Cui, Yuehui He

**Author notes:** These authors contributed equally (Shu Tian, Xiao Luo).

## Abstract

In many flowering plants, the developmental switch from vegetative growth to reproduction (flowering) is timed by seasonal changes in the length of daylight (photoperiod). Under inductive day lengths, the photoperiod pathway typically generates rhythmic expression of a transcriptional activator for florigen production. In the facultative long-day plant *Arabidopsis thaliana*, long-day exposure results in increasing buildup of the CONSTANS (CO) protein towards the end of daylight, and CO activates the expression of the major florigen gene *FLOWERING LOCUS T* (*FT*) to confer long-day induction of flowering. CO-mediated *FT* activation must be properly controlled to prevent an excessive florigen production and precocious flowering under inductive long days, but the underlying mechanism remains elusive. Here, we report an auto-repression mechanism to prevent excessive FT production in inductive photoperiods. We show that the transcription factor FD is expressed in leaf veins and complexes with FT to recognize several *cis*-regulatory DNA motifs in *FT* promoter. FT-FD antagonizes CO-mediated *FT* activation to feedback down-regulate *FT* expression and thus prevent its excessive induction by long-day signals, thereby precluding precocious transition to flowering. Furthermore, we found that in the facultative short-day plant soybean, an *FT* homolog directly represses its own expression. Thus, the auto-repression of *FT* or an *FT* homolog is a conserved mechanism to prevent excessive production of this potent floral regulator in plants, ensuring the floral transition at a proper time to balance vegetative growth with reproductive success and maximize plant production.

## Introduction

In many flowering plants, the developmental transition to flowering or switch from vegetative growth to reproduction phase is timed by seasonal changes of day length through the photoperiod pathway, to achieve reproductive success under favorable seasons (Andres and Coupland, 2012; Song et al., 2015). Light signals are perceived by photoreceptors such as phytochromes and cryptochromes in leaves, and are further integrated with the circadian clock to generate rhythmic expression of a transcriptional activator or regulator for florigen production (Andres and Coupland, 2012; Song et al., 2015). This leads to floral induction under inductive photoperiods.

In the facultative long-day plant *Arabidopsis thaliana*, the *CO-FT* regulatory module determines the photoperiodic flowering response. Under long day (LD) conditions, the circadian-clock regulation of the transcriptional activator *CO* results in a high-level expression from late afternoon into night (Suarez-Lopez et al., 2001; Yanovsky and Kay, 2002). Furthermore, the stability of CO protein is regulated post-translationally. CO is stabilized by far-red and blue light signals, but is destabilized by red light and two RING-finger E3 ubiquitin ligases (Song et al., 2015; Valverde et al., 2004). The coincidence of high-level of *CO* mRNA expression with the presence of both far-red and blue light in late afternoon under LDs, results in the accumulation of CO protein towards the end of daytime (dusk) (Valverde et al., 2004; Yanovsky and Kay, 2002). CO functions to activate the expression of the florigen gene *FT*, the major output of the photoperiod pathway, conferring long-day induction of flowering in Arabidopsis (Andres and Coupland, 2012; Lv et al., 2021).

The CO protein gradually accumulates in leaf veins from late afternoon towards dusk in LDs (Valverde et al., 2004). Upon that it reaches a threshold level with increasing light exposure, the light-stabilized CO protein, together with two NUCLEAR FACTOR-Y (NF-Y) subunits B and C, constitute a trimeric transcriptional activator complex termed as CO-NF (Gnesutta et al., 2017; Lv et al., 2021; Wenkel et al., 2006). This complex recognizes four *CO*-responsive DNA elements (COREs) located in the *FT* proximal promoter to activate *FT* expression in LDs (Tiwari et al., 2010; Zeng et al., 2022). Meanwhile, the distal enhancer bearing the CCAAT motif, recognized by a trimeric NF-Y transcription factor complex, is brought close to the CORE region (bound by CO-NF) by promoter looping, thereby activating *FT* expression (Cao et al., 2014; Liu et al., 2014). *FT* expression is gradually activated from late afternoon and peaks by the end of the day (light period) (Suarez-Lopez et al., 2001; Yanovsky and Kay, 2002). Subsequently, rapid CO degradation by the ubiquitin-proteasome system at night, leads to *FT* repression (Valverde et al., 2004). Hence, the accumulation of CO protein towards the end of daytime results in the rhythmic *FT* activation in LDs, enabling plants to align their timing of the transition to flowering with long-day seasons.

FT, homolog of phosphatidylethanolamine-binding protein (PEBP), functions as a mobile signal and is transported from leaf veins where it is produced to the shoot apical meristem (SAM) through the phloem (Corbesier et al., 2007; Jaeger and Wigge, 2007; Liu et al., 2012; Mathieu et al., 2007). The FT protein in the SAM is received by a 14-3-3 protein (also known as GF14) in the cytoplasm, and FT-GF14 further enters the nucleus to form a trimeric complex with FLOWERING D (FD), a basic Leucine Zipper domain (bZIP) transcription factor (Abe et al., 2005; Taoka et al., 2011; Wigge et al., 2005). This trimeric transcriptional activation complex, termed as the florigen activation complex (FAC), directly activates the expression of the floral meristem identity genes in SAM, including *APETALA1*, *LEAFY* and *CAULIFLOWER* (Abe et al., 2005; Taoka et al., 2011; Wigge et al., 2005). FD has been reported to be expressed preferentially in the SAM to complex with FT to promote the floral transition in Arabidopsis (Abe et al., 2005; Wigge et al., 2005). FT, FD and 14-3-3 proteins are evolutionarily conserved in flowering plants (Li et al., 2015; Taoka et al., 2011). The crystal structure of the rice FAC in complex with DNA has been determined, revealing the direct binding of FD to an ACGT-bearing motif that is facilitated by FT (Taoka et al., 2011).

FT, first discovered in Arabidopsis, is found in all flowering plants examined so far (Andres and Coupland, 2012; Kardailsky et al., 1999; Kobayashi et al., 1999; Putterill and Varkonyi-Gasic, 2016). Multiple *FT* orthologs or homologs exist in various angiosperms, and the functions of *FT-like* genes often have been diversified during recent evolution (Jin et al., 2021; Wickland and Hanzawa, 2015). In the long-day plant sugar beet (*Beta vulgaris*), there are two *FT* paralogs including *BvFT1* and *BvFT2* with opposite function in the regulation of flowering time: *BvFT1* functions as a floral repressor and inhibits the expression of the potent floral inducer *BvFT2* (Pin et al., 2010). In the facultative short-day plant soybean (*Glycine max*), there are 12 *FT* homologs in six homoeologous pairs, among which four *FT-like* genes including *GmFT1a*, *GmFT2a*, *GmFT4* and *GmFT5a* play important roles to regulate flowering (Lin et al., 2021). Under short-day (SD) conditions, both *GmFT2a* and *GmFT5a* are expressed at high levels to strongly promote the floral transition, whereas in LDs both genes are expressed at low levels, but are essential for eventual flowering after prolonged vegetative growth (Cai et al., 2020; Kong et al., 2010). Interestingly, both *GmFT1a* and *GmFT4* are highly expressed in LDs and function to inhibit the floral transition (Liu et al., 2018; Zhai et al., 2014). *GmFT1a* overexpression results in the repression of both *GmFT2a* and *GmFT5a* and consequent late flowering in both SDs and LDs (Liu et al., 2018). It seems that the functionally-antagonistic *FT-like* genes may regulate the expression of their opposing homologs to ensure the appropriate timing of flowering in response to environmental cues, but the underlying molecular mechanisms are unknown.

Plants fine-tune the timing of their transition to flowering to optimize the balance between vegetative growth and reproductive success (Amasino, 2010; Song et al., 2015). A shorter vegetative growth period or early flowering typically results in inadequate accumulation of resources for maximal seed production (Amasino, 2010). On the other hand, a delay in flowering can result in missing the optimal timing and environmental conditions necessary for maximizing seed production in the field, as evidenced by overexpression of a floral inhibitor in rice resulting in a ten-day delay in flowering and a significant reduction in grain yield in the paddy fields (Wang et al., 2021). Because *FT* or *FT* relatives play a central role to promote the transition to flowering in angiosperms (Andres and Coupland, 2012; Song et al., 2015), the proper level of FT production is crucial to balance vegetative growth and reproduction. Under inductive day lengths, the gradual accumulation of CO protein towards the end of light period results in an increasing buildup of CO (Valverde et al., 2004), which may lead to CO over-accumulation. This consequently is expected to give rise to an excessive transcriptional activation of *FT*, resulting in precocious transition to flowering. How day-length induction of FT production is properly controlled to prevent its overproduction, is essentially unknown.

Here we report a negative feedback loop governing *FT* expression to prevent excessive FT production under inductive long days in Arabidopsis. We found that the bZIP transcription factor *FD* is expressed in leaf veins to downregulate *FT* expression and thus delay the floral transition. Upon the CO protein accumulation towards dusk, *FT* expression is strongly activated around dusk. We show that the FT protein complexed with FD (FD-FT) recognizes several *cis*-regulatory motifs in *FT* promoter to antagonize CO-mediated *FT* activation. FD-FT inhibits CO-trigged *FT* promoter looping in transcriptional activation. Our results reveal that FT feedback downregulates its own expression to prevent its excessive induction by the long-day photoperiod pathway, thereby precluding precocious transition to flowering in Arabidopsis. Furthermore, we found that the soybean *FT* homolog *GmFT4*, like *FT*, directly represses its own expression. Thus, the auto-repression of *FT* or an *FT* homolog is a conserved mechanism to prevent excessive production of this potent floral regulator in flowering plants, ensuring the plants to flower at a proper time to balance vegetative growth with reproductive success under a given environmental setting. Moreover, we found that *GmFT4* directly represses the expression of *GmFT2a* and *GmFT5a* to inhibit soybean flowering, revealing a molecular mechanism for the cross-regulation of antagonistic *FT-like* genes in soybean and likely in other flowering plants.

## Results

### *FT* represses its own expression

It is well known that FT partners with the bZIP transcription factor FD to activate the expression of floral meristem identity genes to promote the floral transition (Abe et al., 2005; Wigge et al., 2005), and the FT-FD complex has been thought to function mainly to activate target gene expression. In an effort to elucidate photoperiodic regulation of *FT* expression, we unexpectedly found that in a weak loss-of-function *ft* allele, *ft-1*, bearing a missense *ft* mutation (glycine 171 to glutamate (Kardailsky et al., 1999)), the level of *ft-1* transcript was higher than *FT* in the wild type Col-0 (WT) over a long-day cycle (Fig. 1A). We further examined the stability of *ft-1* and *FT* mRNAs, and found that the *ft-1* transcript degraded moderately faster than the *FT* mRNA (Supplementary Fig.1A). Thus, the increased level of *ft-1* mRNA (relative to the native *FT*) is not attributed to its stability. These results together indicate that *FT* may repress its own expression. To determine whether *FT* is essential for its own downregulation, we introduced into *ft-1* a recombinant *FT* expression reporter gene in which the reporter gene *β-GLUCURONIDASE* (*GUS*) is driven by a 6.9-kb *FT* promoter (Hu et al., 2021). Subsequently, we compared *FTpro-GUS* expression in the leaves of WT and *ft-1* (at ZT12 in LDs), and found that it was apparently increased in this weak *ft* allele (Fig. 1B and Supplementary Fig. 1B and C). Thus, *FT* functions to repress its own transcription.

**Figure 1.**
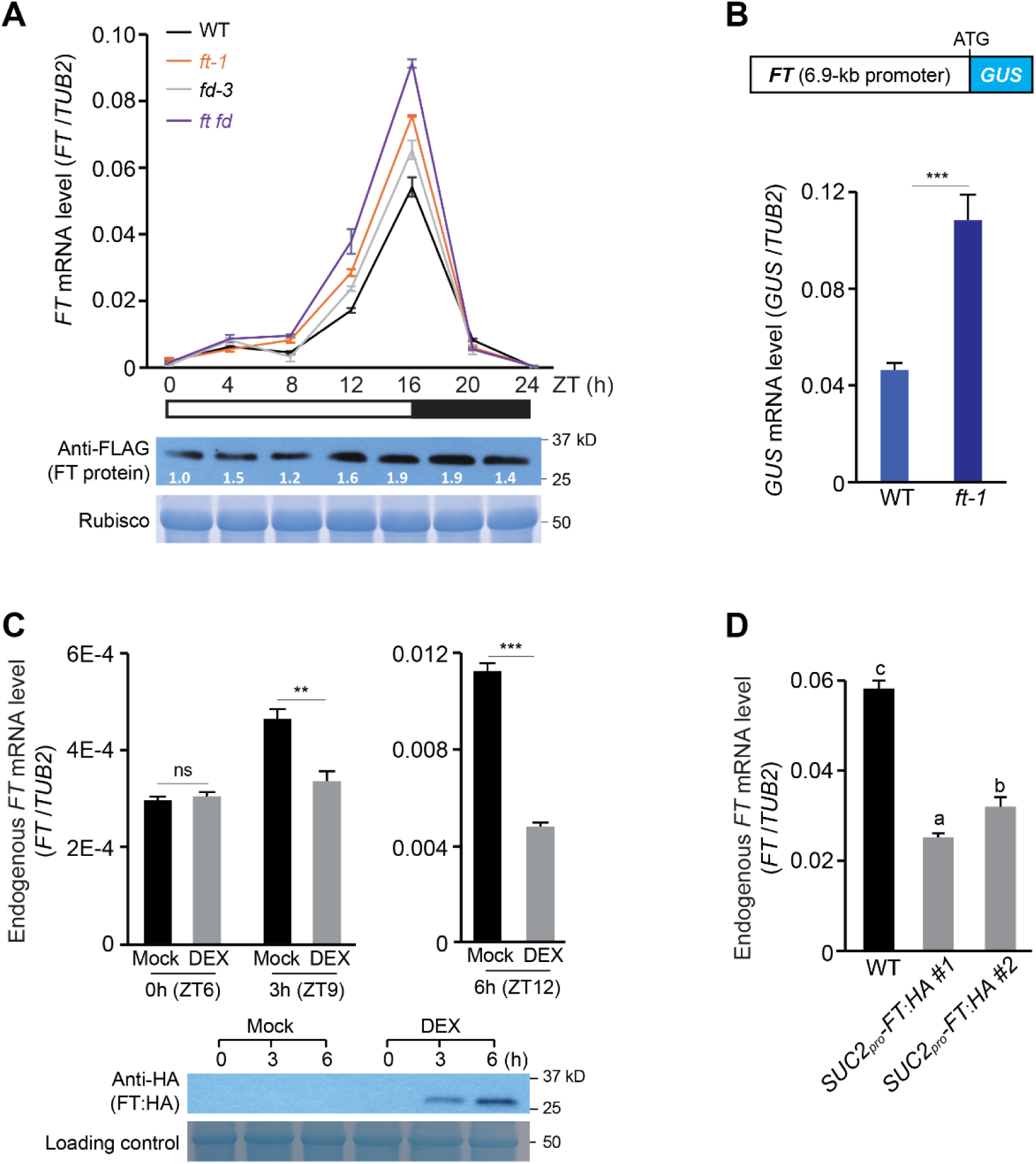
*FT* represses its own expression. **A)** *FT* expression in the rosette leaves of WT (Col-0), *ft-1*, *fd-3* and *ft fd* over a 24-h LD cycle. Levels of *FT* or *ft* transcript were measured by quantitative PCR and normalized to the constitutively-expressed *TUBULIN2* (*TUB2*), and *FT* mRNA expression is shown on the upper panel. Shown on the panel bottom are the levels of FT:FLAG in rosette leaves over a LD cycle, examined by immunoblotting with anti-FLAG (Rubisco serving as a loading control). **B)** *FTpro-GUS* expression in rosette leaves of the indicated T_1_ transgenic plants grown in LDs. Leaves from 25 to 30 independent T_1_ seedlings per line were harvested at ZT12 and pooled for RNA extraction in each sample. **C)** Levels of the endogenous *FT* transcript upon *FT:HA* induction by DEX in the seedlings grown in LDs. Seedlings were treated with DEX (20 mM) at ZT6, and subsequently were harvested at ZT9 and ZT12. The levels of the FT:HA protein induced ectopically by DEX were examined by immunoblotting and shown in the panel bottom, with Rubisco as a loading control. **D)** Endogenous *FT* expression in rosette leaves of WT and *SUC2_pro_-FT:HA* transgenic lines. Samples were harvested at ZT16. **A-D)** The levels of *FT* or *GUS* transcript were normalized to *TUBULIN2* (*TUB2*), and values are means ± standard deviation (s.d.) of three biological replicates. In (**B-C**), two-tailed *t* tests were conducted (** *p*< 0.01, *** *p*< 0.001, and ns for not significant), whereas in (**D**), one-way ANOVA was conducted (letters for statistically-distinct groups; *p*<0.01).

To further confirm that *FT* feedback represses its own expression, we first constructed an *FT* induction line expressing a stringent glucocorticoid-inducible *FT* expression cassette (*FT:HA*, tagged by epitope HA) (Craft et al., 2005). Subsequently, *FT* expression was ectopically induced in seedlings by dexamethasone (DEX). We found that six hour (h) after DEX application, the endogenous *FT* expression was greatly repressed (Fig. 1C). Thus, *FT* indeed represses its own expression.

*FT* is specifically expressed in the phloem tissues in leaf veins (Takada and Goto, 2003). We overexpressed *FT:HA* by the phloem-specific promoter of *SUCROSE TRANSPORTER 2* (*SUC2*) (Truernit and Sauer, 1995). *FT:HA* overexpression in leaf veins resulted in early flowering (Supplementary Fig. 1D), consistent with previous findings (Liu et al., 2012). Next, we examined endogenous *FT* expression in two independent transgenic lines of *SUC2_pro_-FT:HA*, and found that the expression of native *FT* was greatly suppressed in these lines (Fig. 1D). These results further confirm that *FT* represses its own expression in leaf veins.

*FT* mRNA expression is rhythmically regulated under inductive LDs (16-h night/8-h night): strongly activated around dusk (ZT16), but repressed at night and into the early afternoon under controlled environment (Suarez-Lopez et al., 2001; Yanovsky and Kay, 2002). We found that *FT* represses its own expression from late afternoon until dusk (Fig. 1A). Consistently, we found that the FT protein (FT:FLAG) accumulated in leaf veins at a relative high level from late afternoon until night (Fig. 1A), in line with a previous study (Kim et al., 2016). Taken together, these results reveal that upon the induction of *FT* expression by the LD pathway from late afternoon towards dusk, *FT* feedback represses its own expression.

### *FD* is expressed in leaf veins and functions to repress *FT* expression

FT, a PEBP homolog unable to regulate transcription by itself, often complexes with the bZIP transcription factor FD or FD PARALOG (FDP) to regulate target gene expression (Abe et al., 2005; Jaeger et al., 2013; Wigge et al., 2005). Therefore, we measured *FT* mRNA levels in *fd* (Abe et al., 2005) and *fd ft* over a long-day cycle, and found that *FT* expression is de-repressed upon loss of *FD* function (Fig. 1A), consistent with the notion that FT complexes with FD to repress its own expression. In addition, we observed that the level of *FT* de-repression was moderately higher in *ft fd* than either single mutant (Fig. 1A), which may be attributed to that *ft-1* is a weak allele. FD is well known to function as a potent floral activator, and constitutive expression of *FD* by the potent *35S* promoter results in early flowering (Abe et al., 2005; Wigge et al., 2005). It has been shown that *FD* is mainly expressed in SAM where the FD protein functions as part of FAC to promote the floral transition (Abe et al., 2005; Wigge et al., 2005). We reasoned that FD may be expressed in the phloem cells of leaf veins and partners with FT to repress *FT* expression in a feedback manner. To elucidate the role of FD in leaf veins, we examined a transgenic line *FD_pro_-GFP:FD* which can completely rescue the late-flowering phenotype of the *fd* null mutant (Collani et al., 2019). Using cryosectioning of leaf veins of GFP:FD, followed by confocal imaging, we observed that the FD was localized in phloem cells (companion cells) and SAM (Fig. 2A), consistent with the notion that FT-FD represses *FT* expression in leaf veins.

**Figure 2.**
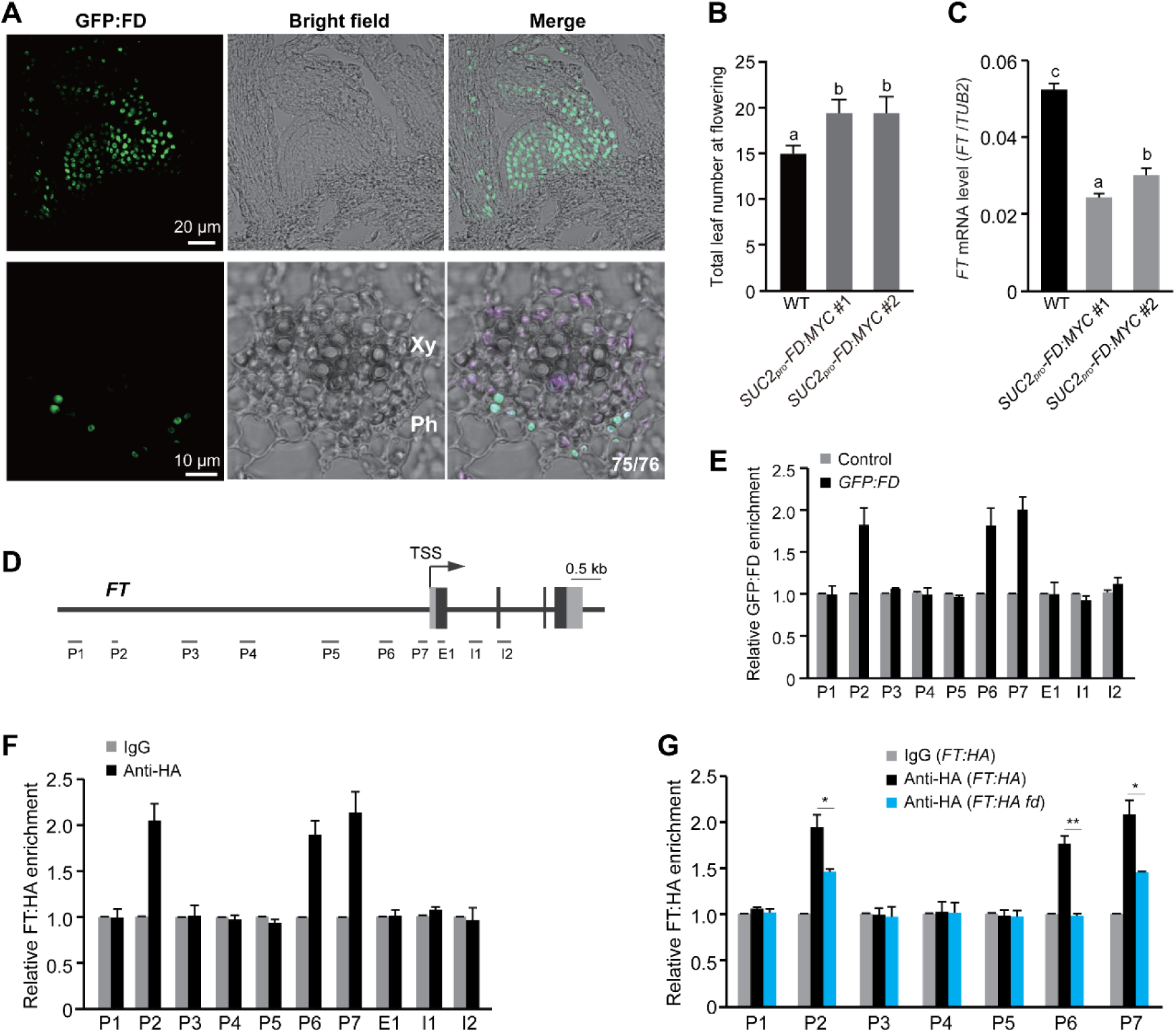
FD is expressed in leaf veins and complexes with the FT protein to down-regulate *FT* expression. **A)** Confocal imaging of the GFP:FD localization in the shoot apical region (longitudinal sections in top panel) and major leaf veins (transverse sections in bottom panel). Ph, phloem; Xy, xylem; purple color for autofluorescence. **B)** Flowering times of *SUC2pro-FD:MYC* lines grown in LDs. Total number of leaves formed prior to flowering was scored (16 plants per genotype). Error bars indicate s.d. Letters indicate statistically-significant differences (one-way ANOVA; *p*<0.01). **C)** Analysis of *FT* expression in rosette leaves of WT and *SUC2pro-FD:MYC* lines grown in LDs. **D)** Schematic diagram of the *FT* gene. Filled gray boxes, untranslated regions (UTRs); filled black boxes, coding regions; arrow, transcription start site (TSS). Chromatin immunoprecipitation (ChIP)-examined regions are indicated with grey bars. **E)** ChIP analysis of GFP:FD enrichment at the *FT* locus. Immunoprecipitated *FT* DNA fragments by anti-GFP were quantified by qPCR and normalized to the internal background gene *TUB2*. Shown are the fold enrichments of GFP:FD at each *FT* region in an *FDpro-GFP:FD* line over background control (WT). **F)** ChIP analysis of FT binding to its own promoter. Total chromatin extracted from the *FT_pro_-FT:HA* seedlings (in *ft-1*) was immunoprecipitated using anti-HA or anti-IgG (background control). Relative FT fold enrichments at each examined *FT* region in the *FT:HA* line over the background control are shown. Notably, the *FT:HA* line carries a single-locus transgene and the endogenous *ft-1*, and anti-IgG serves as a background control for ChIP. **G)** FD is required for FT binding to its own promoter regions, as revealed by ChIP-qPCR with anti-HA. Relative FT:HA fold enrichments in each *FT* promoter regions over the background control anti-IgG are presented. **C** and **E-G)** Samples were harvested at ZT16. Values are means ± s.d. of three biological replicates. Letters in (**C**) indicate statistically-significant differences (one-way ANOVA; *p*<0.01). Two-tailed *t* test was conducted in (**G**), with * *p*< 0.05, and ** *p*< 0.01.

*FT* is specifically expressed in the phloem tissues of leaf veins. To further examine *FT* repression by FD in the phloem tissues, we introduced the transgene of *FD* (fused with the epitope *Myc*) driven by the phloem-specific *SUC2* promoter (*SUC2pro-FD:Myc*) into WT. Subsequently, we examined four independent transgenic lines grown in LDs, and found that all exhibited late flowering (Fig. 2B and Supplementary Fig. 2A and B); furthermore, we found that *FT* expression was greatly repressed in the transgenic lines (Fig. 2C). In addition, we observed that *CO* expression was not affected upon *FD:Myc* overexpression (Supplementary Fig. 2C). Thus, FD functions to repress *FT* expression in leaf veins to delay the transition to flowering. Taken together, these results reveal that FD plays a dual role in the regulation of flowering time: downregulating *FT* expression in leaf veins, but complexing with the FT protein in SAM to promote the floral transition.

### FD-FT directly represses *FT* expression

To explore how FT regulates its own expression, we generated a transgenic line expressing an FT protein (tagged with HA) driven by *FT* promoter (*FT_pro_-FT:HA*), which partly rescued the late flowering phenotype of *ft-1* (Supplementary Fig. 1E). Next, we conducted chromatin immunoprecipitation (ChIP) assays, and found that FT:HA was strongly enriched in one distal region and two proximal regions of *FT* promoter (Fig. 2D and F). Thus, FT directly regulates its own expression.

FT, lacking a DNA-binding domain, complexes with FD to regulate target gene expression. We further conducted ChIP assays using the transgenic seedlings expressing the functional *FD_pro_-GFP:FD*, and found that FD specifically bound the *FT* promoter regions where the FT protein binds (Fig. 2E). This is consistent with that FD complexes with FT to repress *FT* expression in leaf veins.

Next, we determined whether the FD protein is required for FT binding to its own promoter regions. We first introduced *fd* into an *FT:HA* line by genetic crossing to generate an *FT:HA fd* line. Subsequent ChIP assays revealed that the FT enrichments at both distal and proximal FT promoter regions were strongly reduced upon loss of *FD* function (Fig. 2G). Thus, FD indeed is required for FT binding to its own promoter. Previously, it has been shown that in the FAC complex FD directly binds DNA, whereas FT stabilizes the complex and may also contact DNA (Taoka et al., 2011). We further explored that whether FT may be partly required for FD binding FT promoter regions by conducting ChIP with an *GFP:FD ft* line, and found that loss of *FT* function resulted in an apparent reduction of FD enrichment in *FT* promoter regions (Supplementary Fig. 2D). These results, together with that both FT and FD bind the same *FT* promoter regions, reveal that FT complexes with FD to directly regulate its own expression.

### FD recognizes several DNA motifs in *FT* promoter to repress *FT* repression

The bZIP transcription factor FD recognizes ACGT or TCGA-containing motifs in target promoters to regulate gene expression (Abe et al., 2005; Collani et al., 2019; Wigge et al., 2005). We further analyzed the *FT* promoter regions bound by FT-FD, and identified at least one putative FD-binding motif in each region, including C-box (5’-GACGTC-3’), GTCGAC motif and A-box (5’-TACGTA-3’) (Fig. 3A) (Abe et al., 2005; Collani et al., 2019; Taoka et al., 2011). Next, we mutagenized these motifs in the *FT_pro_-GUS* reporter gene, and found that the mutation of each motif resulted in varying degrees of *FT_pro_-GUS* de-repression with the mutation of the C-box motif located in the distal *FT* promoter region exhibiting the greatest effect (Fig. 3B). Furthermore, the mutant *FT_pro_-GUS* (*mFT_pro_-GUS*) in which all three putative FD-bound motifs were mutated exhibited a great de-repression (compared to wild-type *FT_pro_-GUS* in the WT background: over four-fold increase) (Fig. 3C). Taken together, these results show that all three FD-bound motifs are partly required for *FT* repression with the C-box motif in the distal *FT* promoter region playing a major role.

**Figure 3.**
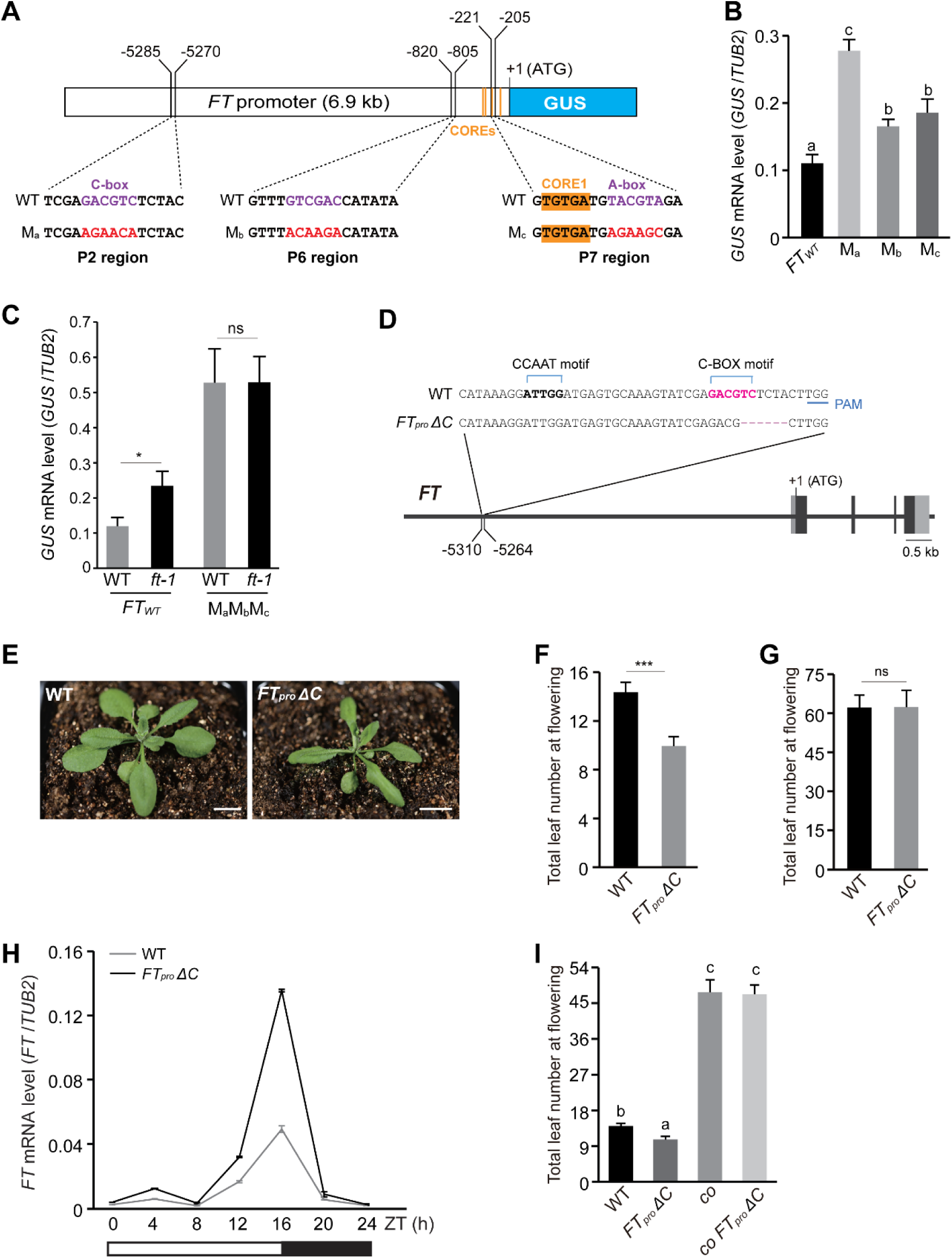
Characterization of the *cis*-regulatory DNA motifs mediating *FT* auto-repression. **A)** Schematic representation of the recombinant *FTpro-GUS* reporter genes consisting of the GUS coding region driven by a 6.9-kb *FT* promoter or a mutated *FT* promoter. M_a_, M_b_ and M_c_ denote mutated FD-binding motifs in *FT* promoter. A of ATG as +1. **B** and **C)** Analysis of *GUS* expression in rosette leaves of M_a_, M_b_, M_c_ (**B**) and M_a_M_b_M_c_ (*mFT_pro_-GUS*) seedlings (**C**) grown in LDs. M_a_, M_b_ and M_c_ are in the WT background. M_a_M_b_M_c_ was introduced into both WT and *ft-1*. Leaves from 25 to 30 independent T_1_ seedlings per line were harvested at ZT16 and pooled for RNA extraction in each sample. Levels of *GUS* transcript were quantified by RT-qPCR and normalized to *TUB2*. Values are means ± s.d. of three biological replicates. **D)** Schematic illustration of CRISPR/Cas9-mediated mutagenesis of the C-box motif in the distal *FT* promoter. *FT_pro_ ΔC* bears a 6-bp deletion, including 2 bp from the C-box. PAM, protospacer adjacent motif. Note that NF-Y recognizes the CCAAT motif to activate *FT* expression. **E)** Early-flowering phenotype of *FT_pro_ ΔC* in LDs. Scale bars, 1 cm. **F** and **G)** Flowering times of WT and *FT_pro_ ΔC* in LDs (**F**) and SDs (**G**). 16-20 plants per genotype were scored, and bars for s.d. **H**) Diurnal expression of *FT* mRNA in rosette leaves of WT and *FT_pro_ ΔC* over a 24-h LD cycle. Levels of *GUS* transcript were directly normalized to *TUB2*. Values are means ± s.d. of three biological replicates. **i**, Flowering times of the indicated lines grown in LDs. Total number of leaves from around 12 plants per line was scored, and bars indicate s.d. **B**, **C**, **F**, **G** and **I)** Two-tailed *t* tests were conducted in (**C**, **F** and **G**), \**p*<0.05, \*\*\**p*<0.001 and ns for not significant; in addition, one-way ANOVA was conducted in (**B** and **I**), and letters mark statistically significant differences (*p*<0.01).

To determine whether the great de-repression of *mFT_pro_-GUS* is attributed to a disruption of *FT* auto-repression, we introduced this transgene into *ft-1*. Subsequent measurements of *GUS* expression revealed that the level of *mFT_pro_-GUS* de-repression in both *ft-1* and WT was nearly identical (Fig. 3C), suggesting that these *cis*-regulatory motifs mediate *FT* auto-repression. Interestingly, we noticed that the extent of *mFT_pro_-GUS* de-repression in *ft-1* was greater than the extent of *FT_pro_-GUS* de-repression in *ft-1* (Fig. 3C). This may be attributed to that the ft-1 protein is still partially functional.

The major motif C-box for *FT* repression is located 18 bp downstream of the essential enhancer element (CCAAT) for CO-mediated *FT* activation (Cao et al., 2014; Liu et al., 2014); hence, it was of great interest to explore the role of this distal C-box in the *FT* repression by FT-FD. Using a CRISPR/Cas9-based method, we generated a homozygous mutant, *FT_pro_ ΔC*, with a six-bp deletion (5’-<u>TC</u>TCTA-3’) that removed two bp of the C-box (5’-GACG<u>TC</u>-3’) (Fig. 3D). Next, we measured *FT* expression in this mutant over a LD cycle, and found that *FT* expression was strongly de-repressed in *FT_pro_ ΔC* compared to WT at ZT16 (Fig. 3H). This confirms that the C-box indeed plays an important role for *FT* repression. The *FT_pro_ ΔC* mutant exhibited early flowering in LDs, but not in SDs (Fig. 3F and G). This long day-dependent early flowering phenotype is consistent with the role of C-box in *FT* auto-repression, namely, the long day-induced CO accumulation is required for *FT* repression. Note that the CO protein is degraded rapidly in SDs. We further introduced a *co* null mutation into *FT_pro_ ΔC*, and found that *co FT_pro_ ΔC* exhibited the same late-flowering phenotype as *co* (Fig. 3I), confirming that the early-flowering of *FT_pro_ ΔC* depends on the *CO* function in LDs. Taken together, these results show that the C-box plays an important role for long day-dependent *FT* auto-repression.

We further assessed the biological function of the C box-mediated *FT* auto-repression by measuring the indicators of overall plant growth and development, including biomass and seed yield (Lim et al., 2018). We found that the early flowering in the *FT_pro_ΔC* line leads to a strong reduction in biomass as well as decreased seed yield compared to WT in LDs (Supplementary Fig. 3A to C). Thus, the C box-mediated *FT* auto-repression generates a proper level of FT production under inductive long days, to balance vegetative growth and reproduction and thus maximize plant production.

### FD binds the C-box in the distal *FT* promoter to inhibit promoter looping

To understand how the regulatory module of FT-FD-C box represses *FT* expression, we first determined whether FD binds the C-box in distal *FT* promoter by ChIP using the GFP:FD and GFP:FD *FT_pro_ ΔC* lines. The enrichment of FD at *FT* promoter was reduced specifically in the C box-containing region upon the partial deletion of C-box, confirming that FD indeed binds this C-box (Fig. 4A). Furthermore, we determined whether C-box is crucial for *FT* repression by FD by introducing *FT_pro_ ΔC* into a late-flowering *SUC2_pro_-FD:MYC* line. We found that the *FT* de-repression in *FT_pro_ ΔC* was largely not restored by *SUC2_pro_-FD*, resulting in early flowering in the *FD:MYC FT_pro_ ΔC* line (Fig. 4B and Supplementary Fig. 3D). In addition, we found that the transcriptional repression of endogenous *FT* by *SUC2_pro_-FT* was relieved by the C-box mutation in *FT_pro_ ΔC* (Supplementary Fig. 3E). These results together show that FT-FD binds the C-box in distal *FT* promoter to mediate *FT* auto-repression.

**Figure 4.**
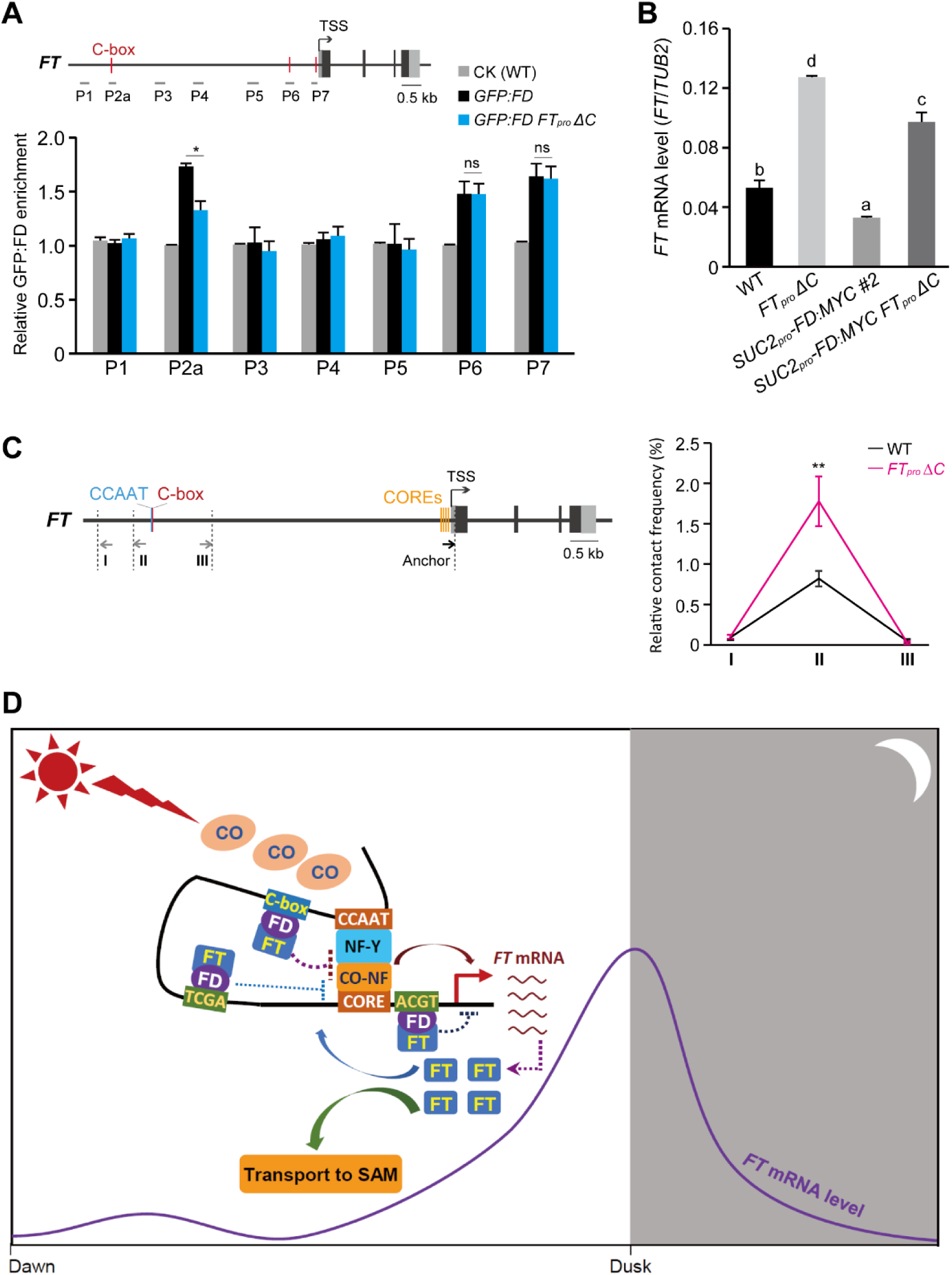
FT-FD binds the C-box in distal *FT* promoter to disrupt *FT* promoter looping and attenuate CO-NF-mediated *FT* activation under inductive LDs. **A)** ChIP analysis GFP:FD binding to *FT* promoter regions upon C-box deletion (ΔC). Shown are relative fold enrichments of GFP:FD across *FT* promoter over a background control (WT). Samples were harvested at ZT16. Values are means ± s.d. of three biological replicates. Two-tailed *t* tests were conducted to evaluate means differences (* *p*< 0.01 and ns for not significant). **B)** *FT* expression in rosette leaves of indicated lines. The levels of *FT* transcript were quantified by RT-qPCR and normalized to *TUB2*. Values are means ± s.d. of three biological replicates. Letters mark statistically significant differences (one-way ANOVA, *p*<0.01). **C)** Quantitative 3C analysis of *FT* promoter looping between the distal CCAAT enhancer and the proximal CORE region in WT and *FT_pro_ ΔC* seedlings grown in LDs. On the left is a schematic diagram of *FT* gene, with vertical dotted lines to denote *Dpn II* cutting sites; the CORE region is set as the anchor, and arrows next to the cutting sites indicate primers. Relative contact frequencies between the anchor and indicated regions were calculated as described in the Methods. Samples were harvested at ZT16. Values are means ± s.d. of three biological replicates, and evaluated by two-tailed *t* test (** *p*<0.01). **D)** A working model for auto-downregulation of the florigen FT production in Arabidopsis under inductive LDs. From late afternoon to dusk, increasing buildup of the CO protein leads to the increased assembly of the transcriptional activator complex CO-NF; in addition, the distal CCAAT enhancer recognized by NF-Y frequently interacts with the proximal CORE region bound by CO-NF, resulting in promoter looping for transcriptional activation. The FT proteins synthesized in leaf veins (companion cells) complex with FD, and the FD-FT complex binding the distal C-box (next to the CCAAT enhancer), downregulates the looping frequency of *FT* promoter; in addition, FD-FTs binding the A-box within the CORE region and the G<u>TCGA</u>C motif close to CORE also act to attenuate CO-NF-mediated *FT* activation. In short, there is a ‘braking’ mechanism ensuring a proper level of FT production to balance vegetative growth with reproduction success for improved biomass and seed yield under inductive photoperiods.

*FT* promoter looping between the distal CCAAT-bearing enhancer region and the proximal CO-bound region with CO-responsive elements (COREs) is essential for CO activation of *FT* expression in inductive LDs (Cao et al., 2014; Liu et al., 2014). The FT-FD-bound C-box is next to CCAAT. We speculated that the FT-FD-C-box regulatory module may interfere with CO-triggered *FT* promoter looping to downregulate its expression. Hence, we measured the interaction frequencies between the proximal CORE region and distal regions at the *FT* promoter in WT and *FT_pro_ ΔC*, using the chromosome conformation capture (3C) approach. We found that the looping frequency between the CCAAT enhancer region and the CORE region was greatly increased upon the C-box removal (Fig. 4C). Thus, the FT-FD-C-box regulatory module indeed inhibits the interaction of the distal CCAAT enhancer with the proximal promoter region to downregulate CO-mediated *FT* activation in inductive LDs.

Our results thus far show that FD-FT binds *FT* promoter, particularly the distal C-box, to antagonize CO-mediated *FT* activation and thus feedback downregulate *FT* expression (Fig. 4D). This prevents an excessive *FT* induction by the long-day photoperiod pathway in Arabidopsis.

### An *FT-like* gene represses its own expression and other *FT-like* genes in soybean

To elucidate whether *FT* auto-repression is a conserved mechanism to regulate *FT-like* gene expression in flowering plants, we explored the regulation of *FT-like* genes in soybean, a facultative short-day plant with multiple *FT-like* genes (Lin et al., 2021). In response to SDs, both *GmFT2a* and *GmFT5a* are expressed at relative high levels to promote flowering, whereas under LDs, *GmFT1a* and *GmFT4* are highly expressed to inhibit the floral transition, resulting in late flowering (Lin et al., 2021). We generated several transgenic lines in which *GmFT4* (tagged with *FLAG*) was overexpressed (*GmFT4^OE^:FLAG*) (Fig. 5A and Supplementary Fig. 4A). We found that in both SDs and LDs, *GmFT4* overexpression strongly repressed the expression of the endogenous *GmFT4* (Supplementary Fig. 4B and D). Next, we conducted ChIP assays with *GmFT4^OE^:FLAG* seedlings, and found that GmFT4 was enriched at three *GmFT4* promoter regions bearing putative FD-binding motifs (ACGT-containing elements) (Fig. 5C). Note that FD is evolutionarily conserved in soybean (Lin et al., 2021). Thus, like FT-FD in Arabidopsis, GmFT4 directly represses its own expression in soybean. This reveals the auto-repression of *FT* or an *FT* homolog is a conserved mechanism to prevent its excessive production.

**Figure 5.**
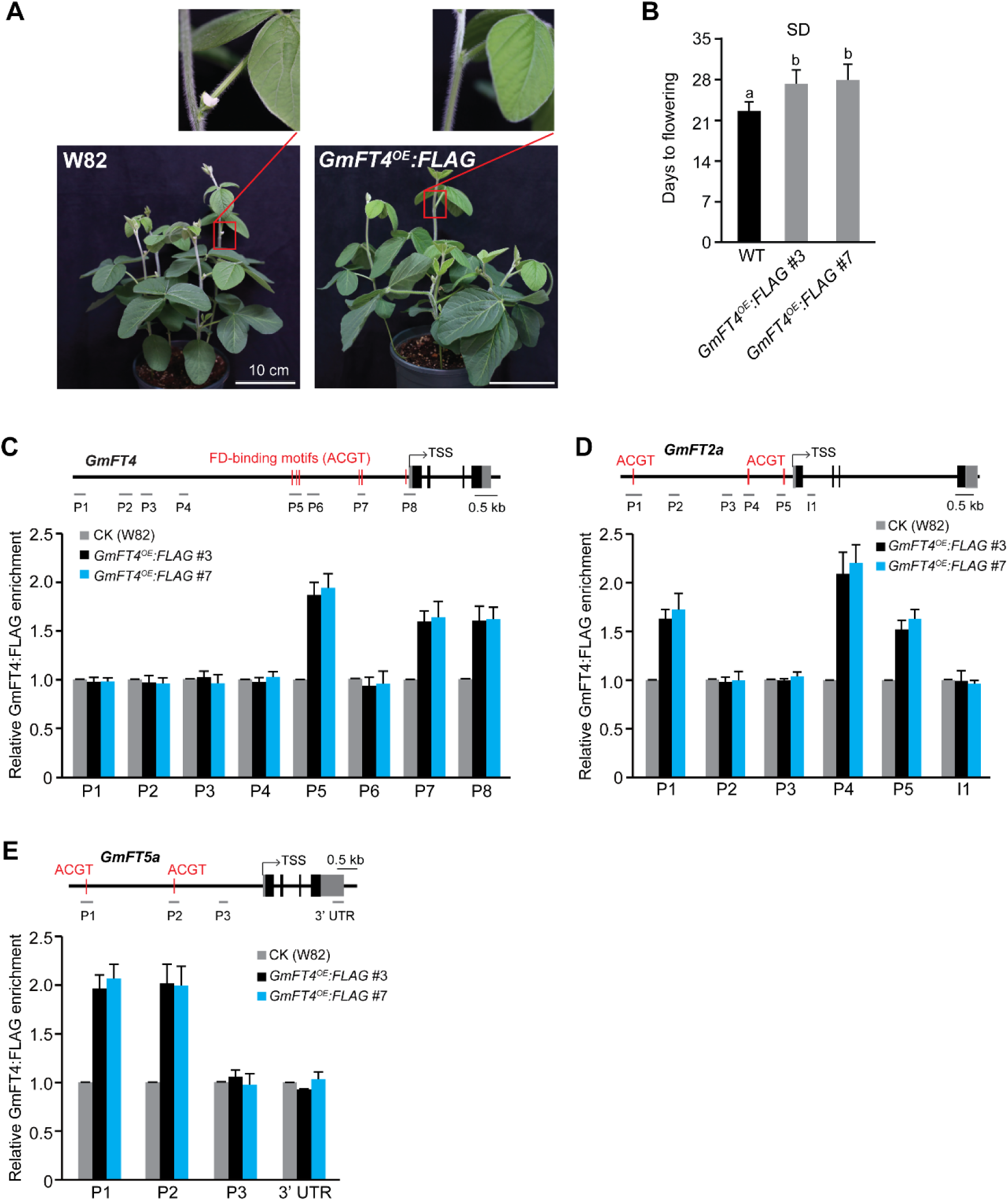
*GmFT4* directly represses its own expression as well as *FT-like* genes in soybean. **A)** Phenotypes of the transgenic lines overexpressing *GmFT4:FLAG* (*GmFT4^OE^:FLAG*) grown for 21 days after cotyledon emergence (DAE) in SDs. Scale bars, 10 cm. **B)** Flowering times of *GmFT4^OE^:FLAG* lines grown in SDs. About 17 plants for each line were scored. Kruskal-Wallis nonparametric analysis with Dunn’s multiple comparison tests was conducted, and letters indicate statistically significant differences (*p*<0.01). **C-E)** ChIP-qPCR analysis of GmFT4 enrichment at the *GmFT4* (**C**), *GmFT2a* (**D**) and *GmFT5a* loci (**E**). On top of each panel is a schematic drawing of a soybean *FT-like* gene (filled gray boxes for UTRs, and filled black boxes for coding regions), with red lines to denote putative FD-binding motifs and grey bars for ChIP-examined regions. Total chromatin was extracted from the leaves of two independent *GmFT4^OE^:FLAG* lines grown in SDs (harvested at ZT8), followed by immunoprecipitation with anti-FLAG. DNA fragments were quantified by qPCR and normalized to the internal control *Glycine max ACTIN 11* (*GmACT11*). Shown are relative fold enrichments of GmFT4:FLAG over background control (the non-transgenic background *Williams 82*). Values are means ± s.d. of three biological replicates.

Functionally antagonistic *FT-like* genes have been widely found in flowering plants, the floral repressors of *GmFT1a* and *GmFT4* may repress the expression of *GmFT2a* and *GmFT5a* to inhibit flowering in LDs. Recently, it has been shown that the relative expression levels between *GmFT4* and *GmFT2a*/*GmFT5a* play an important role to determine soybean flowering under different day lengths (Lee et al., 2021). We found that *GmFT4* overexpression represses both *GmFT2a* and *GmFT5a* expression, but not *GmFT1a* expression (Supplementary Fig. 4C, E and F), resulting in a delay in flowering under SDs (Fig. 5B). Furthermore, we uncovered that GmFT4:FLAG was enriched at multiple *GmFT2a* and *GmFT5a* promoter regions bearing FD-binding motifs (with the ACGT core) (Fig. 5D and E), revealing a direct regulation by GmFT4. These results are consistent with the notion that under LDs the highly-expressed GmFT4 directly represses the expression of both *GmFT2a* and *GmFT5a* to inhibit flowering in soybean. Thus, in a flowering plant with multiple *FT* paralogs or homologs, an FT-like protein may not only downregulate its own expression, but also directly modulate the expression of its homologs to determine an appropriate timing to flower in response to day length signals.

## Discussion

In this study, we have found that a negative feedback loop regulates the expression of the florigen *FT* to prevent excessive FT production and precocious flowering under inductive photoperiods. We show that the bZIP transcription factor FD is expressed in leaf veins and that FT-FD specifically binds several ACGT/TCGA-containing motifs in *FT* promoter to downregulate or attenuate CO-mediated *FT* activation from late afternoon to dusk when the CO protein is built up with increasing light period in inductive LDs. We further show that FT-FD functions to inhibit CO-triggered *FT* promoter looping in transcriptional activation. Thus, FT-FD antagonizes the transcriptional *FT* activation by CO-NF, leading to a downregulation of *FT* expression. Together, these findings define an auto-repression mechanism to prevent excessive FT production upon the accumulation of the photoperiod pathway output (CO), conferring an appropriate timing to flower in response to an inductive long-day signal. Furthermore, we have found that in the facultative short-plant soybean, the *FT* homolog *GmFT4*, like *FT* in Arabidopsis, directly represses its own expression. Thus, the auto-repression of *FT* or an *FT* homolog is a conserved mechanism to prevent excessive production of this potent floral regulator in flowering plants.

The coincidence of the circadian-regulated *CO* transcription and exposure to light (blue and far-red light) gives rise to an increasing buildup of the CO protein towards the end of daytime under inductive long-day conditions (Andres and Coupland, 2012; Song et al., 2015). The fluctuating light conditions (both light quality and period) in a given growth period are expected to lead to an over- or under-accumulation of the CO protein, which may over- or under-drive transcriptional activation of *FT* expression. The auto-repression of *FT* provides an important layer of regulation to buffer the fluctuations in CO-mediated *FT* activation. This ensures that the florigen FT is produced at an appropriate level to balance vegetative growth with reproduction under a particular season, as evidenced by that a disruption of auto-repression of *FT* results in a great reduction in biomass and seed yield under a controlled environmental setting.

*FT*, encoding a PEBP protein, is evolutionarily conserved in flowering plants (Jin et al., 2021; Putterill and Varkonyi-Gasic, 2016). The *PEBP* gene family in angiosperms consists of three clades including *MOTHER OF FT AND TFL1* (*MFT*)- *like*, *TERMINAL FLOWER 1* (*TFL1*)*-like* and *FT-like*. *TFL1-like* genes typically function as floral repressors, whereas *FT-like* genes often act as floral inducers (Jin et al., 2021; Wickland and Hanzawa, 2015). During recent evolution of *FT-like* genes through gene duplication and evolutionary innovation in flowering plants, the *FT-like* clade has generated antagonistic regulators that function to fine-tune the timing of floral induction and other developmental transitions, in response to environmental signals (Jin et al., 2021; Wickland and Hanzawa, 2015). For instance, the two *FT* paralogs in sugar beet function oppositely to regulate flowering with *BvFT1* repressing *BvFT2* expression (Pin et al., 2010). In the biennial crop onion (*Allium cepa L.*), there are four *FT* paralogs, *AcFT2* and *AcFT1* promote the floral transition and bulb formation in LDs, respectively, whereas *AcFT1* expression is repressed by *AcFT4* to prevent bulb formation in SDs (Lee et al., 2013). In the SD plant potato (*Solanum tuberosum*), the *FT-like* gene *SELF PRUNING 5G* (*StSP5G*) represses the expression of the *FT* ortholog *StSP6A* in LDs to prevent tuberization (Abelenda et al., 2016). In soybean, long-day photoperiods lead to a high-level expression of *GmFT1a* and *GmFT4*, which repress the expression of the floral activators *GmFT2a* and *GmFT5a*, resulting in an inhibition of the floral transition (Lin et al., 2021). Therefore, it is common in flowering plants that an *FT-like* gene represses the expression of its homologs to fine-tune developmental transitions. In this study, we have found that *GmFT4* directly represses the expression of *GmFT2a* and *GmFT5a* to inhibit soybean flowering, pointing a molecular mechanism for the cross-regulation of antagonistic *FT-like* genes in soybean and likely other flowering plants.

It has been long known that FD is expressed preferentially in SAM and functions as a potent inducer of the floral transition (Abe et al., 2005; Wigge et al., 2005). In this study, we have uncovered that FD is expressed in leaf veins and complexes with the FT protein to repress *FT* expression and thus delay the floral transition under inductive LDs. Hence, FD plays a dual role in long-day induction of flowering: highly expressed in the SAM and complexed with FT to activate floral initiation, and moderately expressed in leaf veins and complexed with FT to prevent FT overproduction in inductive photoperiods, ensuring an appropriate photoperiodic flowering response. In addition to FD, FD homologs may be involved in *FT* autorepression. We found that the FT enrichments at two *FT* promoter regions are partly dependent on the FD protein, suggesting FD homologs such as FDP may be involved in recruiting FT to target regions, as evidenced by that FDP partners with FT to promote the floral transition in SAM (Jaeger et al., 2013). FD is evolutionarily conserved in flowering plants (Li et al., 2015; Lin et al., 2021). In soybean, FD-like proteins complex with FT-like proteins to regulate the expression of flowering-regulatory genes (Lin et al., 2021). We have found that GmFT4 is enriched specifically in the regions bearing FD-binding motifs (with the ACGT core) in *GmFT4* promoter, suggesting that GmFT4 complexes with a FD-like protein to repress its own expression. These findings collectively reveal that the FT-FD complex can function as a transcriptional repressor to form an autoregulatory loop that mediates the feedback repression of *FT* or *FT-like* genes in flowering plants. Noteworthily, the rice FAC formed in leaf veins can feedback activate *Hd3a* expression indirectly (Brambilla et al., 2017), suggesting that multiple regulatory loops are involved in transcriptional regulation of *FT* or *FT-like* genes to fine-tune flowering time, in response to environmental cues.

In summary, we have uncovered an auto-repression mechanism to prevent excessive production of the florigen FT in Arabidopsis and likely in other flowering plants under inductive photoperiods. This results in an appropriate level of the mobile FT or FT-like proteins generated in leaf veins, to optimize vegetative growth for improved seed yield or reproductive success by preventing precocious transition to flowering in response to photoperiodic induction.

## Materials and Methods

### Plant materials and growth condition

All Arabidopsis mutants and transgenic lines used in this study are in the Columbia (Col-0) background. *ft-1*, *ft-10*, *fd-3* and the transgenic line *FD_pro_-GFP:FD* have been described previously (Abe et al., 2005; Collani et al., 2019; Kardailsky et al., 1999; Yoo et al., 2005). Seeds were germinated on agar plates with half-strength MS media, and plants were grown at around 22°C in LDs (16-hour light/8-hour dark) or SDs (8-hour light/16-hour dark) with cool white light.

The soybean cultivar *Williams 82* was used in this study. Plants were grown at around 26°C under LDs (16-hour light /8-hour dark) or SDs (12-hour light/12-hour dark).

### Plasmid construction

The previously described *FT_pro_*-*GUS* fragment(Hu et al., 2021) was cloned into the binary vector *pBGW*(Karimi et al., 2005) via Gateway technology (Invitrogen). To generate *FT_pro_-FT:FLAG* and *FT_pro_-FT:HA*, a 11.0-kb genomic fragment of *FT* (8.9 kb upstream of ATG plus the 2.1-kb genomic coding sequence except for the stop codon) was fused in frame with three copies of FLAG or one copy of HA at 3’ end, respectively, and the fusions were cloned into *pHGW* (Karimi et al., 2005) using Gateway technology. To create *SUC2_pro_-FT:HA* and *SUC2_pro_-FD:MYC*, the 2.0-kb *AtSUC2* promoter was first fused with 0.5-kb *FT* or 0.8-kb *FD* coding sequence (without the stop codon) tagged with three copies of *HA* or four copies of *MYC* (at 3’ end), respectively; subsequently, the fusions were cloned into the binary vector *pBGW* via Gateway technology.

To construct the sgRNA*^FT^*-CRISPR/Cas9 expression cassette, an sgRNA fragment targeting an *FT* promoter region was introduced into *psgR-Cas9-At* at *Bbs I* (Liu et al., 2015), and then the *sgRNA*-*Cas9* cassette was cloned into the binary vector *pCAMBIA1300* at *Hind III* and *EcoR I*. For *35S_pro_-GmFT4:FLAG* construction, the 0.5-kb coding sequence of *GmFT4* (*Glyma.08G363100*) from the reference cultivar *Williams 82* was fused in-frame with three copies of FLAG at 3’ end; subsequently, the fusion was cloned into the binary vector *pB2GW7* (Karimi et al., 2005) using Gateway technology.

### Gene expression analysis

Total RNAs were extracted from leaves or seedlings grown in LDs, with *Eastep Super Total RNA* extraction kit (Promega) according to the manufacturer’s instructions. DNA digestion and reverse transcription were conducted using *HiScript III 1st Strand cDNA Synthesis kit* (+gDNA wiper) according to the manufacturer’s instructions (Vazyme); subsequently, real-time quantitative PCR (qPCR) was performed on an *ABI QuantStudio5 Flex* real-time PCR system using a SYBR qPCR master mix (Vazyme). Transcript levels of gene of interest were normalized to a constitutively-expressed reference gene in Arabidopsis or soybean. Primers are listed in Supplementary Table 1.

### Yeast two-hybrid assay

Yeast two-hybrid assays were conducted according to the instructions from *Matchmaker GAL4 Two-Hybrid System* (Clontech). Firstly, the coding sequences of *CLF* and *FT* were cloned into *pGADT7* and *pGBKT7*, respectively. Paired plasmids were introduced into the yeast strain AH109; subsequently, the transformed yeast cells were grown on a solid synthetic medium without leucine (L), tryptophan (W), histidine (H), and adenine (A) to exam potential protein-protein interaction.

### ChIP assay

ChIP experiments in Arabidopsis were conducted following previously established protocols (Wang et al., 2014), with minor changes. In brief, total chromatin was extracted from the seedlings of interest grown in LDs, followed by immunoprecipitation with anti-HA (Sigma, H9658), anti-FLAG (Sigma, M8823), or anti-GFP (Abcam, ab290). The immunoprecipitated genomic fragments of interest were quantified by qPCR on an *ABI QuantStudio5 Flex* real-time PCR system (Applied Biosystems) with iTaq SYBR Green supermix (Biorad). The levels of examined *FT* fragments were normalized to the internal background control *TUBULIN2* (*TUB2*); subsequently, relative fold enrichments were calculated over a background control. Three biological replicates were conducted for each ChIP assay.

ChIP assays in soybean were carried out as previously described with minor modifications (Saleh et al., 2008). Briefly, approximately 1.0 g of fully expanded trifoliate leaves from 14-d-old soybean plants grown in SDs was collected at ZT8, and fixed with 1% formaldehyde with vacuum infiltration. Subsequently, the nuclei were isolated following an established protocol (Louwers et al., 2009), followed by total chromatin extraction and subsequent fragmentation by Bioruptor (Diagenode). Immunoprecipitations were carried out with anti-FLAG beads (Sigma, M8823). Genomic DNA fragments were quantified by qPCR, and subsequently normalized to the internal background control *GmACT11*. Relative fold enrichments were calculated over a background control. Values are means ± s.d. of three biological replicates. Primer sequences are described in Supplementary Table 1.

### Chromosome conformation capture (3C) experiment

3C assays were conducted as previously described with minor modifications (Cao et al., 2014; Louwers et al., 2009; Luo et al., 2018). Briefly, nuclei were extracted from about 2-g seedlings cross-linked in 2% formaldehyde, and lysed by 0.15% SDS. Next, chromatin-bound DNA was digested by *Dpn II* (NEB) at 37°C overnight, followed by ligation with T4 DNA ligase (NEB) at 16°C for 6 h. After reversing crosslinks, the ligated DNA was purified by phenol-chloroform extraction, followed by ethanol precipitation.

Quantitative PCR was conducted to determine relative interaction frequencies between paired regions. The CORE-bearing region near the transcription start site serves as the anchor segment, and a primer from this region is paired with various primers located at distal *FT* promoter regions for the qPCR analysis. An *FT* region lacking a Dpn II cutting site was used as a loading control, to normalize the differences in DNA concentrations from different samples. Additionally, primer efficiencies were determined using a control template that contained equal amounts of all possible ligation products derived from a 9.1-kb *FT* fragment that had been digested with Dpn II. Relative contact or interaction frequencies were calculated by normalizing the level of an amplified fragment from WT or *FTproΔC* over that from the control template. Primers are listed in Supplementary Table 1.

### Histological β-Glucuronidase activity staining

GUS activity staining was conducted as previously described (Tao et al., 2017). Briefly, transgenic seedlings (T_1_) grown in LDs were harvested at ZT12, and stained in a 1.5 mM X-Gluc (5-bromo-4-chloro-3-indolyl-β-D-glucuronic acid) solution at 37℃ for 12 h, followed by de-chlorophylling with 70% ethanol. Stained seedlings were examined under a Zeiss dissecting microscope.

### Biomass and seed yield assay

The whole plant biomass assay was conducted as previously described with minor adjustments (Lim et al., 2018). In brief, plants were cultivated in soil until a main stem of 1 cm in height had developed after bolting. Fresh weights of 10 seedlings for each line were measured, followed by a full dehydration at 65°C for 24 hours and dry weight measurement.

Seed yields per plant were measured as follow. Plants were grown in soil until no flowers were produced, followed by withholding water for approximately 4 weeks to ensure complete drying of the inflorescences. Subsequently, seeds were harvested from individual plants, and thoroughly dried, and the total dry seed mass per plant was measured.

### Cryosectioning and confocal imaging

Samples were cryo-sectioned and examined with a confocal microscope as described previously with minor modifications (Liu et al., 2019). Briefly, leaves or shoot apical regions from seedlings grown in LDs, were fixed in 4% paraformaldehyde (in Phosphate-Buffered Saline /PBS, pH 6.9) at room temperature for 1 h after vacuum infiltration. Next, the samples were washed by PBS and incubated in 30% sucrose (in PBS) overnight, followed by embedding in Tissue Tek O.C.T. compound (Sakura Finetek) and sectioned to 20-μm thickness by a Leica CM 1950 sliding microtome at - 20℃. Slides were dried in a 42℃ oven and treated briefly with the 1:1 methanol/acetone mixture, and then were mounted with PBS and imaged under a Zeiss LSM900 confocal microscope.

### Soybean transformation

The recombinant *35S_pro_-GmFT4:FLAG* was introduced into the *Agrobacterium tumefaciens* strain *EHA105* and subsequently transformed the soybean cultivar *Williams 82* using the cotyledonary node method as previously described, utilizing the *bar* (bialaphos resistance) gene as a selectable marker in plants with glufosinate as the selection agent (Zeng et al., 2004). Briefly, soybean seeds were surface-sterilized in a desiccator, by exposure to chlorine gas produced from a mixture of hydrochloric acid and sodium hypochlorite. Following overnight sterilization, the seeds were placed on a germination medium plate (24°C for 1 day). After seed germination, the apical shoot/bud was removed, and 3 to 5 incisions (0.5 mm deep and 3-4 mm long) were made at the junction of cotyledon with hypocotyl. The explants were then inoculated with a freshly-prepared *EHA105* culture, followed by co-cultivation, shoot induction and growth, and rooting. Subsequently, the bar-resistant seedlings were transplanted into soil, and the seeds from verified transgenic plants were harvested.

### Statistical analysis

Two-tailed *Student’s t* test and Analysis of One-Way Variance (ANOVA) with the Holm-Sidak method were conducted using GraphPad Prism (v9.5.1). All discrete leaf number data were log-transformed for statistical analysis.

## Acknowledgements

We would like to thank Dingwei He for experimental assistance. This work was supported in part by the National Key R&D Program of China, the National Natural Science Foundation of China, and the Peking-Tsinghua Center for Life Sciences.

## Conflict of interest

The authors declare that they have no conflict of interest.

---

**Supplementary Fig. 1.**
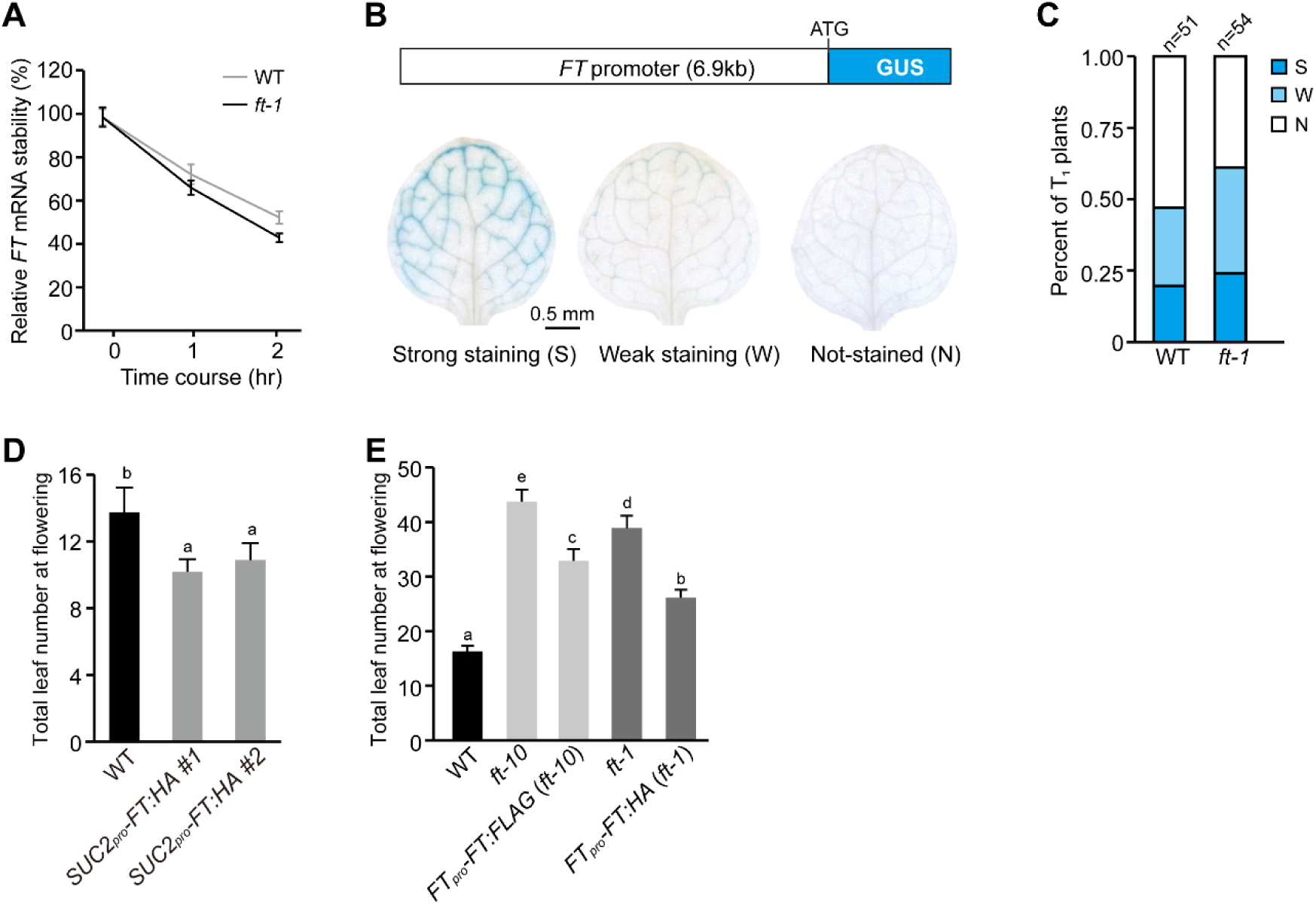
Characterization of auto-repression of *FT* expression. **A)** *FT* mRNA decay in WT and *ft-1* seedlings in LDs. Seedlings were treated with 0.6 mM cordycepin in an incubation buffer (1 mM PIPES pH 6.25, 1 mM sodium citrate, 1 mM potassium chloride and 15 mM sucrose). Transcript levels were quantified by qPCR and first normalized to the internal control 18S rRNA, and relative levels to the initial time point were calculated. Values are means ± s.d. of three biological replicates. **B)** Classification of GUS staining in transgenic seedlings (T_1_ generation) expressing *FTpro-GUS* (in WT) under LD conditions. Seedlings were classified into three groups based on the staining intensity in leaves: strongly stained (S), weakly stained (W), and not stained (N). **C)** Analysis of GUS staining in leaves of the indicated *FTpro-GUS* plants (T_1_ generation) grown in LDs. The percentages of seedlings exhibiting different levels of staining were calculated (n for total number of plants scored in each line). **D** and **E)** Flowering times of *SUC2pro-FT:HA* lines (**D**) and *ft* rescue lines (**E**) grown in LDs. Total number of leaves formed prior to flowering was scored (15-18 plants per line), and bars for s.d. Letters indicate statistically significant differences (one-way ANOVA; *p*<0.01).

**Supplementary Fig. 2.**
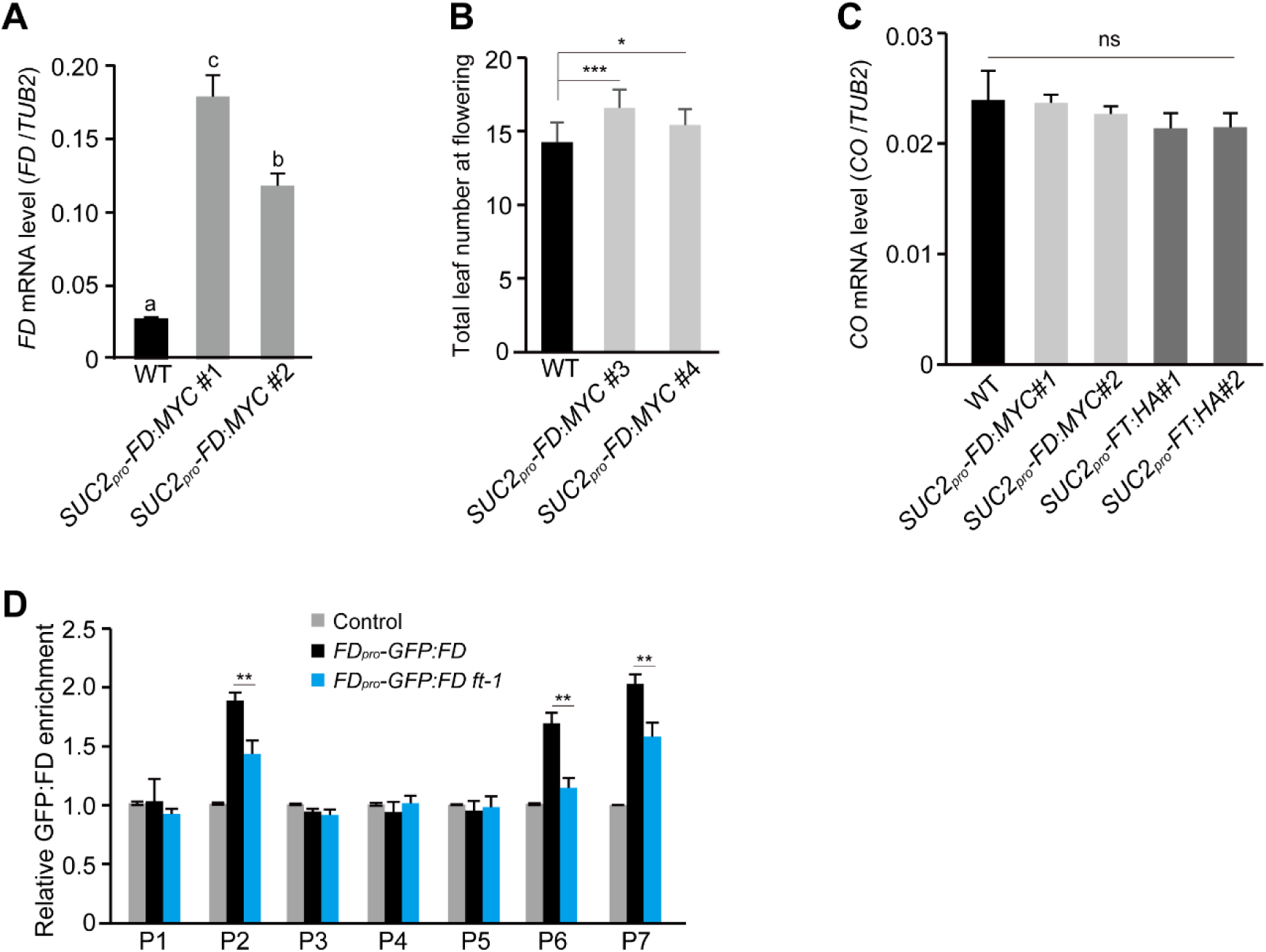
Characterization of *FD-*mediated *FT* repression. **A)** *FD* mRNA expression in rosette leaves of WT and *SUC2_pro_-FD:MYC* lines grown in LDs. Values are means ± s.d. of three biological replicates. **B)** Flowering times of *SUC2pro-FD:MYC* lines grown in LDs. Total number of leaves formed prior to flowering was scored (12 plants per genotype), and bars indicate s.d. **C)** *CO* expression in rosette leaves of WT, *SUC2_pro_-FT:HA* lines and *SUC2_pro_-FD:MYC* lines under LDs (at ZT16). *CO* transcripts were quantified by RT-qPCR and normalized to *TUB2*. Values are means ± s.d. of three biological replicates. **D)** FT is partly required for GFP:FD binding to *FT* promoter regions, as revealed by ChIP-qPCR with anti-GFP. Relative GFP:FD fold enrichments in each *FT* promoter region over the background control (WT IP with anti-GFP) are presented. Samples were harvested at ZT16 (LDs). Values are means ± s.d. of three biological replicates. **A**-**D)** One-way ANOVA was conducted in (**A** and **C**), with letters to indicate statistically significant differences (ns for not significant), whereas in (**B** and **D**) two-tailed *t* tests were carried out (* *p*<0.05, ** *p*<0.01 and *** *p*<0.001).

**Supplementary Fig. 3.**
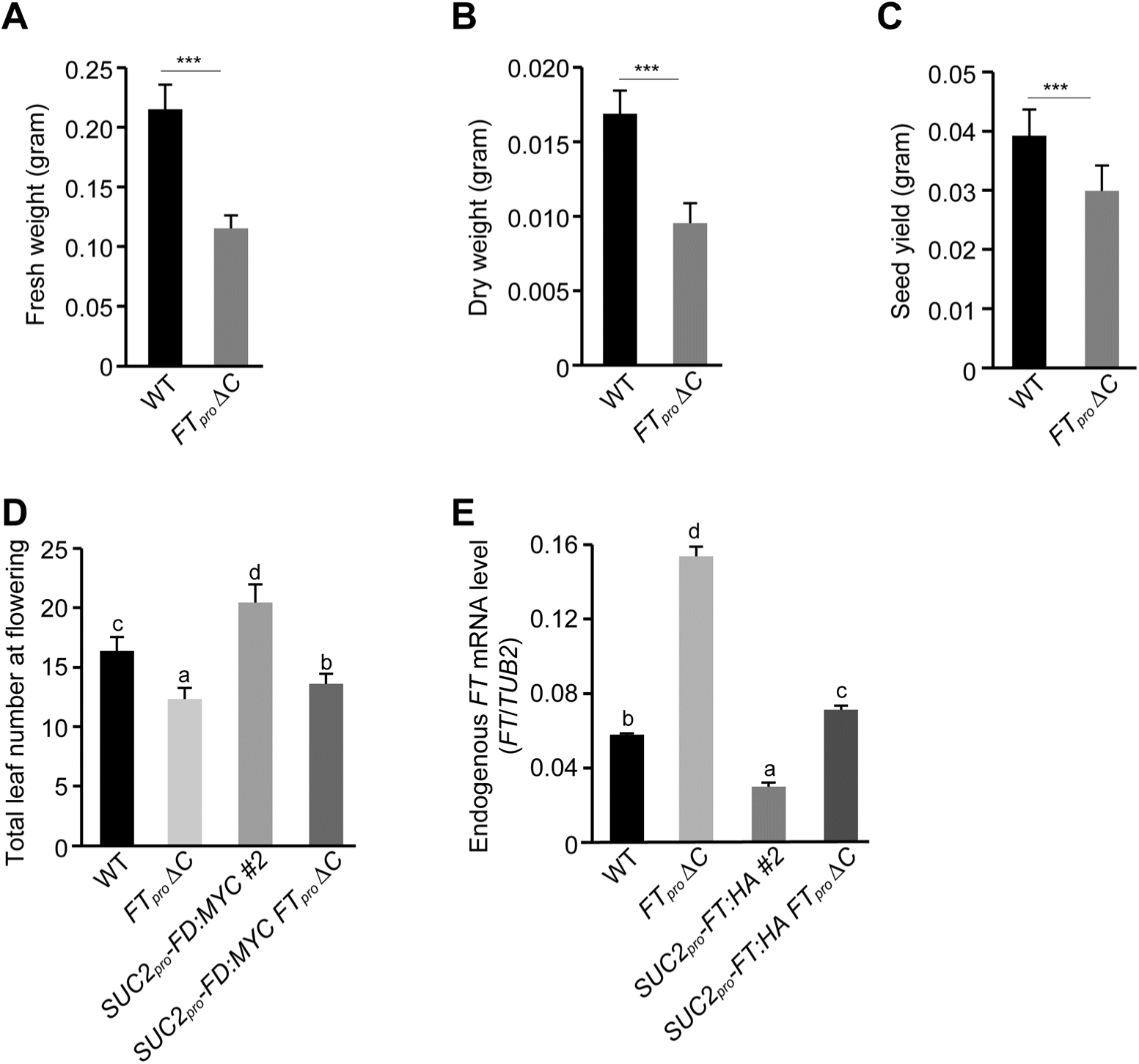
Characterizations of *FT_pro_ ΔC*. **A**-**C**) *FT_pro_ ΔC* exhibits a great decrease in biomass under normal growth conditions in LDs. Fresh weight (**A**), dry weight (**B**) and seed yield (**C**) are measured. A total of 10 plants were scored for each line. Values are means ± s.d. Two-tailed *t* tests were conducted to evaluate statistically significant differences (*** *p*<0.001). **D)** FD-mediated delay in flowering is largely suppressed by *FTpro ΔC*. Total number of leaves formed prior to flowering was scored for each line (30 plants per line grown in LDs). Bars for s.d. and letters denote statistically-significant differences (one-way ANOVA, *p*<0.01). **E)** Downregulation of the endogenous *FT* expression in *SUC2_pro_-FT:HA* lines is partly rescued by *FT_pro_ ΔC*. Seedlings were grown in LDs and harvested at ZT16.

**Supplementary Fig. 4.**
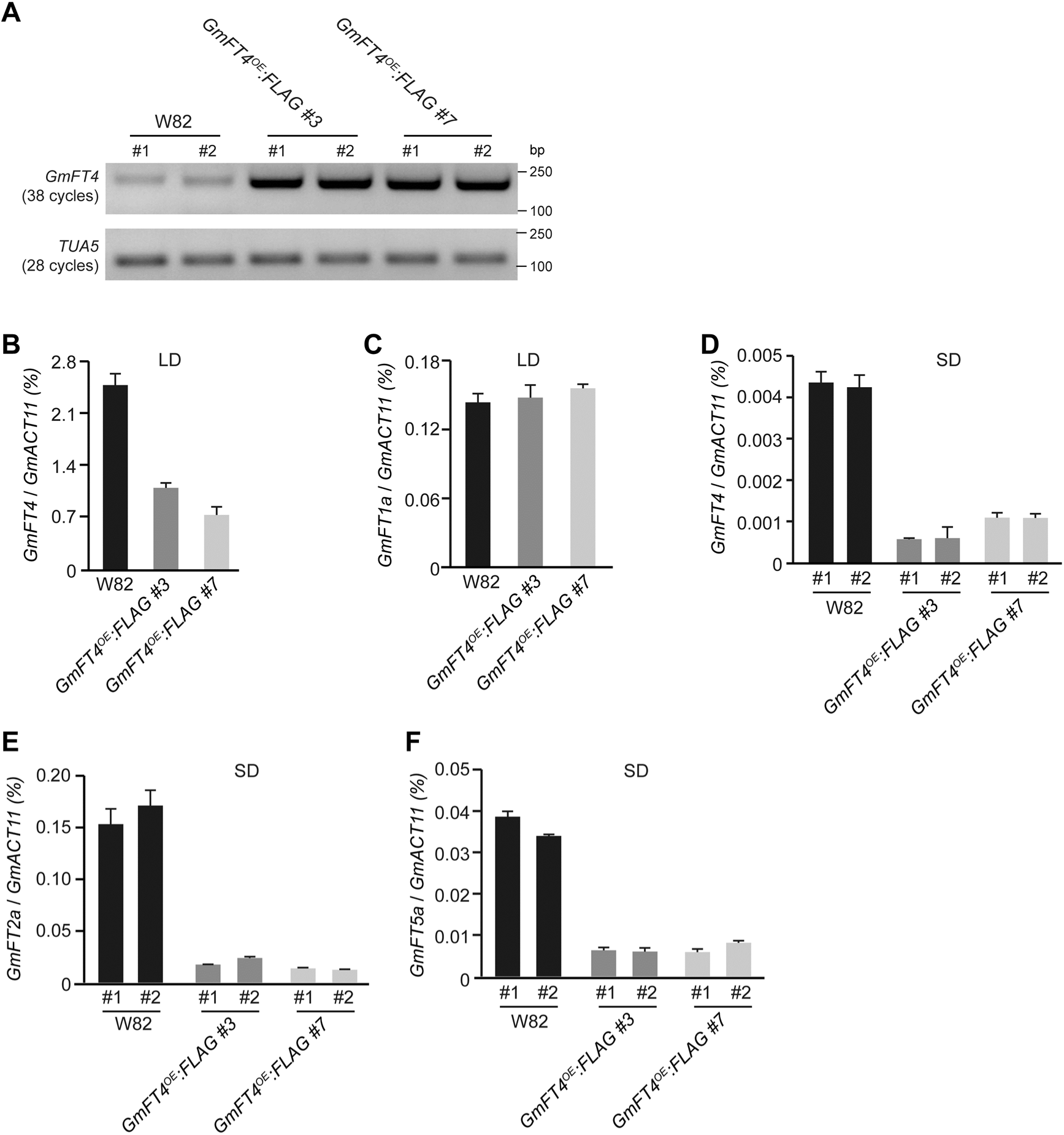
Characterization of *GmFT4* overexpression in soybean. **A)** Analysis of *GmFT4:FLAG* overexpression in leaves of the indicated transgenic soybean lines by semiquantitative RT-PCR. Two individual soybean plants from each transgenic line or *Williams 82* (grown in SDs) were examined. The constitutively-expressed *TUBULIN ALPHA5* (*TUA5*) serves as an internal control. W82 for *Williams 82*, the non-transgenic background. **B** and **C)** Transcript levels of endogenous *GmFT4* (**B**) and *GmFT1a* (**C**) in the leaves of W82 and *GmFT4^OE^:FLAG* lines at 21 DAE under LDs. Leaves from two independent T_3_ soybean lines (5 plants per line) were harvested at ZT16 for RNA extraction. Transcript levels were quantified by RT-qPCR and normalized to the constitutively expressed *GmACT11*. Values are means ± s.d. of three biological replicates. **D-F)** Transcript levels of endogenous *GmFT4* (**D**), *GmFT2a* (**E**), and *GmFT5a* (**F**), and in the leaves of W82 and *GmFT4^OE^:FLAG* lines at 14 DAE under SDs. Samples were harvested at ZT8. Two individual soybean plants from each transgenic line or W82 were examined, and values are means ± s.d. of three technical replicates.

**Supplementary Table 1.**
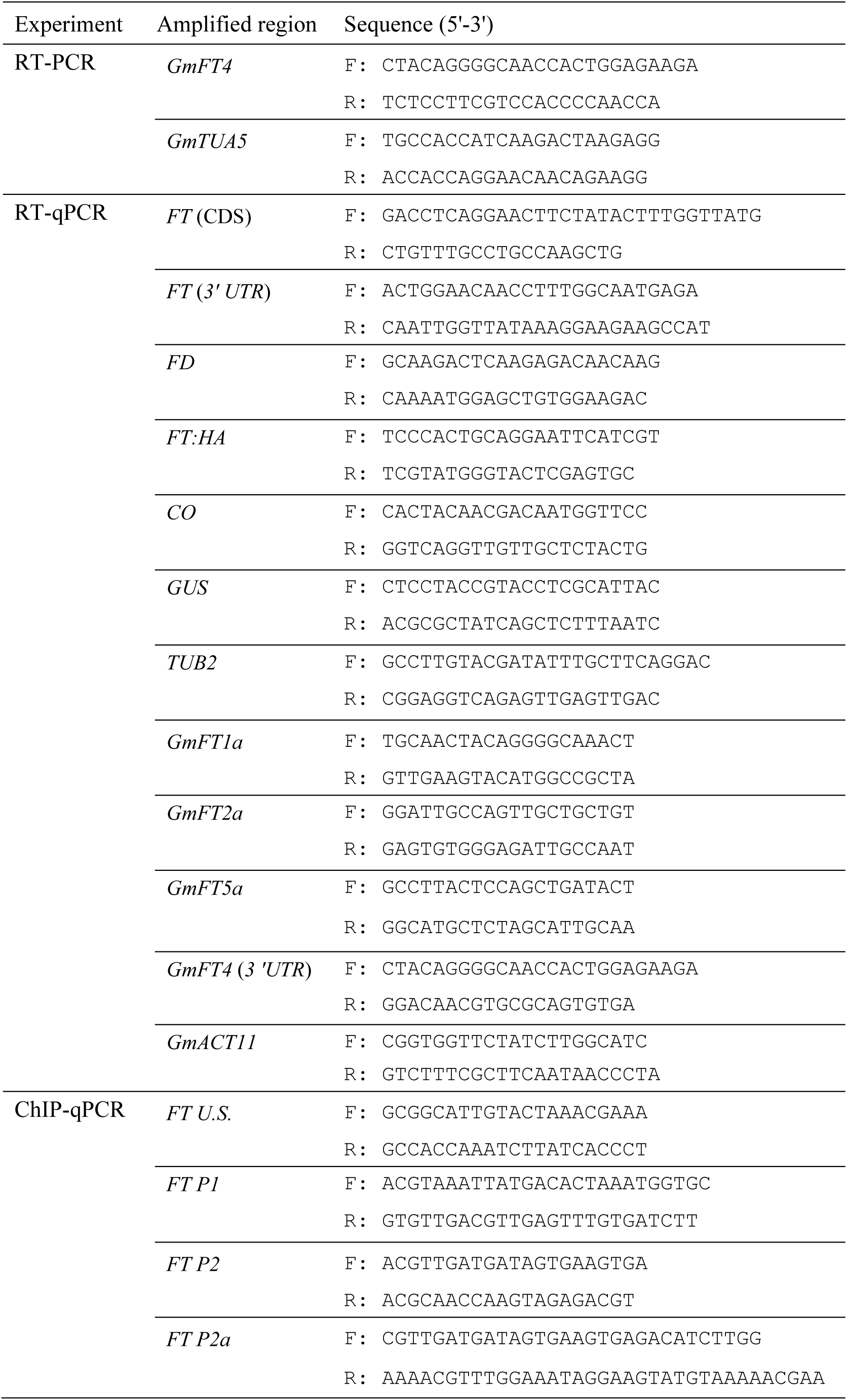
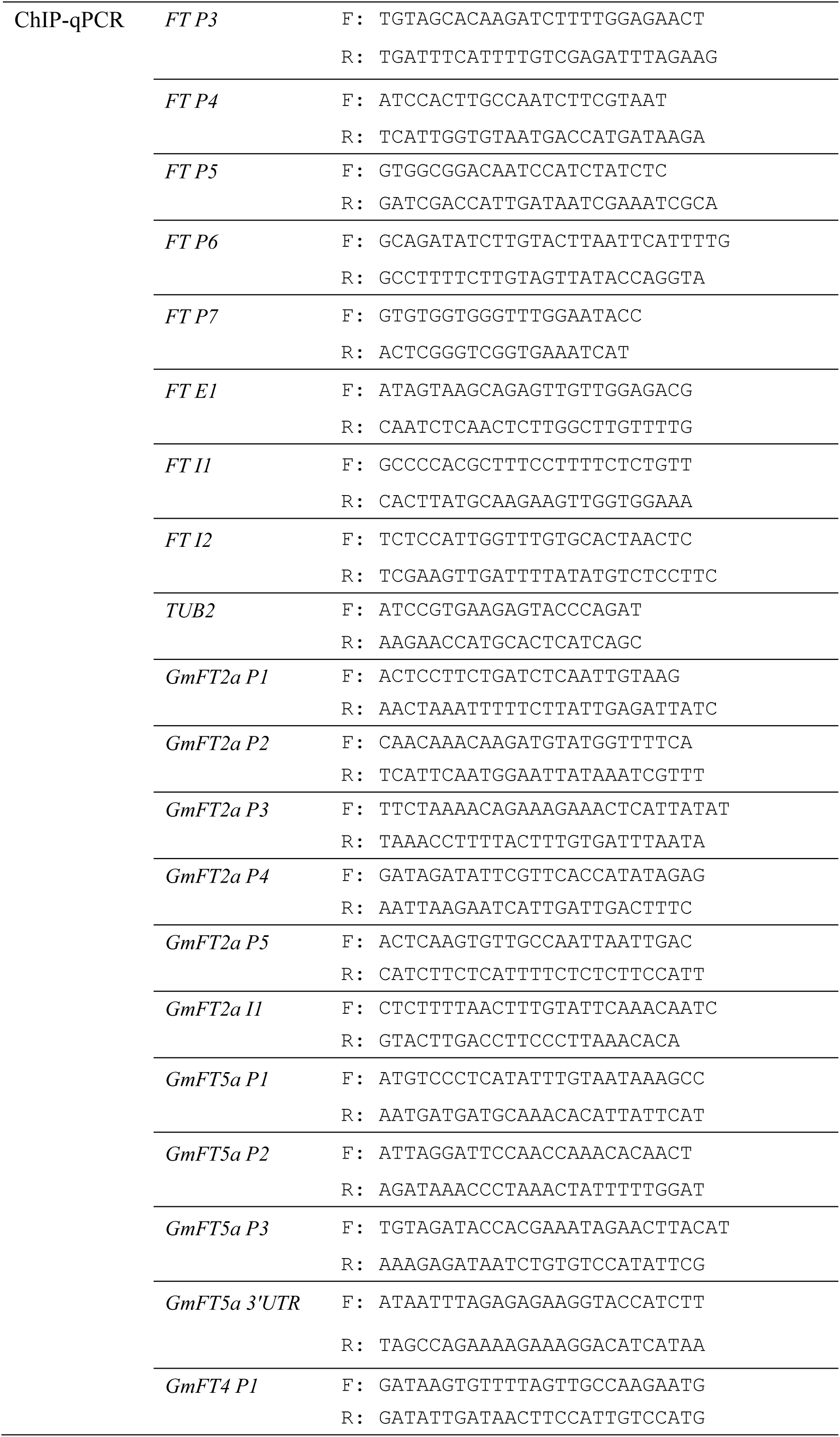
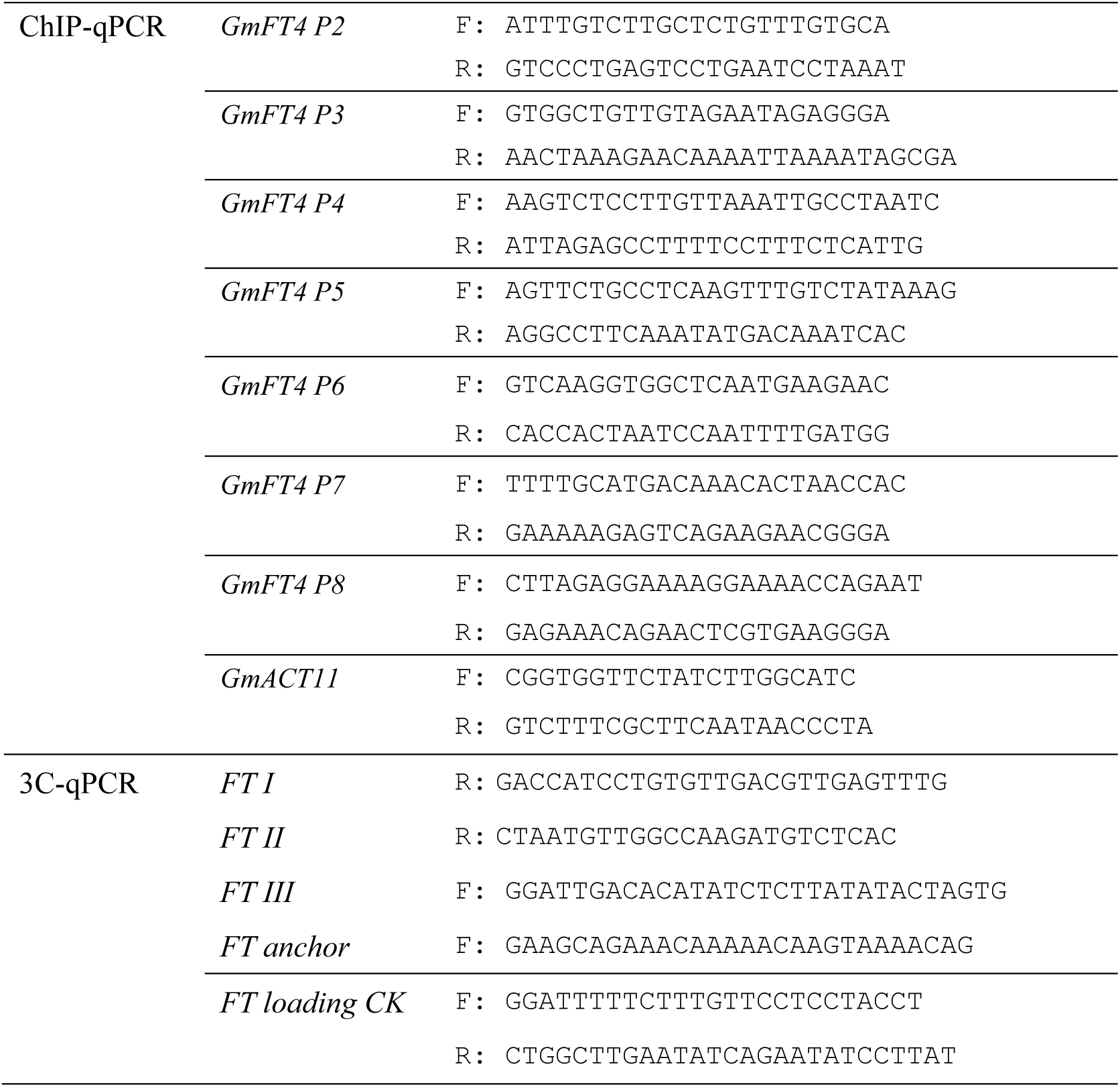
List of primers used in this study.

## Notes

### Competing Interest Statement

The authors have declared no competing interest.

